# A multimodal iPSC platform for cystic fibrosis drug testing

**DOI:** 10.1101/2021.06.21.448578

**Authors:** Andrew Berical, Rhianna E. Lee, Junjie Lu, Mary Lou Beermann, Jake A. LeSeur, Aditya Mithal, Dylan Thomas, Nicole Ranallo, Megan Peasley, Alex Stuffer, Jan Harrington, Kevin Coote, Killian Hurley, Paul McNally, Gustavo Mostovslavsky, John Mahoney, Scott H. Randell, Finn J. Hawkins

## Abstract

Cystic fibrosis (CF) is a monogenic lung disease caused by dysfunction of the cystic fibrosis transmembrane regulator (CFTR) anion channel, resulting in significant morbidity and mortality. The progress in elucidating the role of the CFTR channel using established animal and cell-based models led to the recent discovery of effective CFTR modulators for most individuals with CF. However, a subset of individuals with CF do not respond to these modulators and there is an urgent need to develop novel therapeutic strategies. In this study, we assembled a panel of iPSCs derived from individuals with common or rare variants representative of three distinct classes of CFTR dysfunction. To measure CFTR function in patient-specific iPSCs we adapted two established *in vitro* assays of CFTR function to iPSC-derived airway cells. In both a 3-D spheroid assay using forskolin-induced swelling as well as planar cultures composed of polarized mucociliary airway epithelial cells, we quantified CFTR baseline function and response to CFTR modulators and detected genotype-specific differences. Our results demonstrate the potential of the human iPSC platform as a research tool to study cystic fibrosis and in particular accelerate therapeutic development for CF caused by rare mutations.

## Introduction

Cystic fibrosis (CF) is caused by mutations in the Cystic Fibrosis Transmembrane Regulator (*CFTR*) gene and leads to multi-organ disease particularly affecting the respiratory system. The *CFTR* gene encodes an anion channel involved in the regulation of proper airway surface liquid hydration, viscosity, and pH (Ratjen et al., 2015). Mutations in *CFTR* lead to abnormally viscous secretions in the airways and pancreatic ducts which cause sino-pulmonary infection, bronchiectasis, pancreatic insufficiency, and a shortened life span (Riordan et al., 1989). More than 2000 variants in the *CFTR* gene have been identified to date, with several hundred known to cause clinical disease. *CFTR* mutations are classified according to the molecular defects they cause, including variation in protein quantity (classes 1,5), trafficking (class 2), channel opening (class 3), and ion conductance (class 4) (Cutting, 2014). Due to the range and combinations of *CFTR* mutations, targeted pharmacotherapy approaches are complex. Dependent on their *CFTR* mutation, approximately 90% of individuals with CF should benefit from a newly developed class of medications termed CFTR modulators which improve protein trafficking, function, and patient-centered clinical outcomes (Accurso et al., 2010; Davies et al., 2018; Keating et al., 2018; Middleton et al., 2019; Ramsey et al., 2011; Wainwright et al., 2015). However, CFTR modulators are not curative, expensive, required in combination therapy, and theoretically are required life-long. Furthermore, a subset of individuals with CF caused by class 1 mutations have severe disease and require urgent and ambitious approaches to ameliorate their disease.

Preclinical *in vitro* models were critical to the discovery and approval of CFTR modulators and will almost certainly play a central role in advancing therapeutic options for CF further. Many cellular models of CF exist, including heterologous cell lines, primary human rectal organoids, human nasal epithelial cells and human bronchial epithelial cells (HBECs), each with its own advantages and disadvantages that have been thoroughly reviewed elsewhere (Clancy et al., 2019). On one end of the spectrum, heterologous cell lines transduced with *CFTR* (e.g Fischer Rat Thyroid cells) have enabled high-throughput screening approaches that led to the identification of CFTR modulators but bear little resemblance to the human tissues affected by CF (Sheppard et al., 1994). On the other end of the spectrum are HBECs: the current gold standard cell-based platform for the preclinical assessment of CFTR-mediated ion transport. Obtained from either bronchoscopic biopsy or explanted lungs at the time of transplant, HBECs from individuals with CF differentiate in air-liquid interface (ALI) culture into a pseudostratified airway epithelium with morphologic, molecular and physiologic similarities to the *in vivo* human airway (Fulcher & Randell, 2013; Fulcher et al., 2005). Electrophysiologic studies (eg: Ussing chamber) of CFTR-dependent ion-flux in CF HBEC ALI cultures are sensitive and predictive of *in vivo* response to modulators (Clancy et al., 2019). The efficacy of a candidate drug is typically validated in HBECs prior to advancing to clinical trials. From this *in vitro* pipeline, there are now several FDA-approved CFTR modulators. VX-770, a potentiator that increases the probability of channel opening was the first to be approved for the class 3 gating mutation, Gly551Asp (Ramsey et al., 2011). Clinical treatment with VX-770 leads to a 10% FEV_1_ increase, decreased exacerbations, amongst other patient-centered outcomes. The CFTR corrector molecules VX-809 and VX-661, each used in combination with VX-770 were approved for patients harboring two copies of Phe508del, the most common CF causing CFTR mutation, though the clinical efficacy has been modest (2.6-4% FEV_1_ increase) (Davies et al., 2018; Rowe et al., 2017; Wainwright et al., 2015). A second-generation corrector (VX-445) was recently approved for both homozygous and heterozygous Phe508del patients and shows promising clinical improvement in combination with VX-661 and VX-770 in a large randomized clinical trial (14% FEV_1_ increase) (Middleton et al., 2019). Additionally, data from the preclinical platforms recently formed the basis to expand the approval of CFTR modulators to patients with rarer CFTR mutations not represented in clinical trials (Durmowicz et al., 2018).

While the clinical response to CFTR modulators is encouraging, currently approved drugs do not afford a benefit to ∼10% of all CF patients due to their underlying *CFTR* mutation. Most notably, patients with class 1 non-sense mutations (eg: G542X, W1282X) have severe CF without targeted therapeutic options (Howard et al., 1996; Zainal Abidin et al., 2017). Compared to the class 2 (Phe508del) and 3 (Gly551Asp) mutations above, class 1 mutations are fundamentally different; premature termination codons within mRNA transcripts cause mRNA degradation by nonsense-mediated decay (NMD) or cause premature translation termination, leading to truncated non-functional or absent protein. A successful therapeutic strategy for these individuals will likely involve complex pharmacotherapy, gene-editing, gene-delivery, or cell-based therapy approaches. From a pharmacotherapy approach, combinatorial regimens using read-through molecules (e.g. G418), inhibitors of nonsense-mediated decay (e.g. SMG1i) and modulators are being developed (Keenan et al., 2019; Valley et al., 2019). The development of these ambitious approaches would likely be accelerated by unrestricted access to patient-specific airway cells, however there is a major bottleneck in obtaining HBECs from individuals with rare class 1 *CFTR* variants and electrophysiologic measures of CFTR-dependent current are relatively low throughput. As a result, other *in vitro* platforms and CFTR functional assays are being developed to overcome these hurdles. For example, rectal organoids are quite readily obtained via a minimally invasive biopsy from CF patients and can generate a near limitless supply of patient-specific cells (Clancy et al., 2019; Dekkers et al., 2016; 2013). After rectal biopsy, CFTR-expressing rectal organoids are cultured and CFTR function is measured using the forskolin induced swelling (FIS) assay. Activation of the CFTR channel is initiated by adding forskolin (± CFTR modulator) which leads to a CFTR-dependent increase in organoid size within hours. This platform detected CFTR modulator effects in cells from individuals with different *CFTR* mutations and correlated with *in vivo* drug effects (Berkers et al., 2019; Dekkers et al., 2013; 2016). However, the extent to which a rectal epithelial platform reflects CF disease pathology or is suitable as a readout for class 1 mutations is unknown.

Human induced pluripotent stem cells (iPSCs) have several properties that make them promising candidates for advancing CF therapeutics. iPSCs contain the complete and unique genetic code of the donor, including specific *CFTR* sequence. They are routinely generated by the “reprogramming” of patient-derived somatic cells (eg: dermal fibroblasts, PBMCs) by overexpressing *OCT3/4, SOX2, KLF4*, and *C-MYC* ((Takahashi et al., 2006; Takahashi et al., 2007). By recapitulating key embryological events, iPSCs can be differentiated into alveolar and airway epithelial cells (Chen et al., 2017; Dye et al., 2015; Firth et al., 2014; Hawkins et al., 2021; 2017; S. X. L. Huang et al., 2013; Jacob et al., 2019; Konishi et al., 2016; McCauley et al., 2017; Miller et al., 2018; Mou et al., 2012; Wong et al., 2012). We previously applied these protocols for proof-of-concept CF disease modeling; for example, we corrected the *CFTR* sequence in Phe508del iPSCs and measured CFTR-dependent airway as well as intestinal epithelial spheroid swelling in FIS assays (McCauley et al., 2017; Mithal et al., 2020). Recently, we published the successful derivation of airway basal cells, the major stem cell of the airway epithelium, from iPSCs. These iPSC-derived airway basal cells (iBCs) self-renew *in vitro* and, under similar ALI culture conditions used for HBECs, differentiate into a mucociliary epithelium that is morphologically and transcriptionally similar to primary HBEC ALI cultures (Hawkins et al., 2021). iBC-derived ALI cultures from non-CF individuals expressed *CFTR* and in Ussing chambers exhibited CFTR-mediated transepithelial ion flow at a magnitude expected of non-CF HBECs.

In the present study, we hypothesized that iPSC-derived CFTR expressing airway cells from individuals with class 1-3 CFTR mutations can be used to measure baseline CFTR function as well as rescue in response to CFTR modulator treatment. Two established CFTR functional assays were adapted: (1) FIS of 3-D airway epithelial cell spheroids and (2) electrophysiologic measurement of CFTR dependent current across mucociliary ALI cultures, both of which detect responses to clinically approved CFTR modulators. Furthermore, combinatorial treatment of iPSC-derived airway cells from an individual with a class 1 mutation led to a significant increase in CFTR function. This iPSC-based platform can overcome a major limitation in the development of therapeutic strategies for CF, particularly for individuals with rare mutations, by providing a near limitless supply of patient-specific cells with the ability to quantify CFTR function.

## Results

### CF iPSCs efficiently differentiate into airway epithelial spheroids

In order to develop a standardized approach for the study of multiple classes of CFTR mutations, we first sought to establish a bank of iPSCs from individuals with CF. We identified and reprogrammed five iPSC lines from individuals with class 1-3 mutations in the *CFTR* gene (Fig. 1a). Reprogramming of iPSCs from somatic cells was performed as previously described (Supplementary Fig. 1) (Gianotti-Sommer et al., 2008; Somers et al., 2010; Sommer et al., 2009; Takahashi et al., 2006; Takahashi et al., 2007). We preferentially focused on identifying individuals with homozygous mutations to exclude the potential confounding effect of compound heterozygous mutations from different classes that might obscure the interpretation of CFTR modulator responses. For class 1, iPSCs were previously generated from an individual homozygous for a non-sense mutation in exon 23 – c.3846G>A, p.Trp1282X, was identified (iPSC line “P20801” hereafter referred to as W1282X, provided by Prof. Scott Diamond, University of Pennsylvania). For class 2 mutations, a total of three iPSC lines were studied. Two lines were homozygous for the Phe508del mutation (c.1521_1523delCTT, p.Phe508del), one of which was newly reprogrammed (iPSC line: “CFTR4-2”, hereafter Phe508del #1), and the other was previously described (iPSC line: “RC2 204”, for simplicity referred to hereafter as Phe508del #2). A third iPSC line derived from an individual with compound heterozygous mutations Phe508del/Ile507del (p.Ile507del, c.1519_1521delATC), (iPSC clone “C17”, hereafter Phe508del #3) (see Methods for cell line details) (Crane et al., 2015; Somers et al., 2010). For class 3 mutations, we identified one of the few documented individuals homozygous for the Gly551Asp mutation (c.1652G>A, p.Gly551Asp), and generated an iPSC line (hereafter G551D). We also selected three established non-CF PSC lines as controls (“RUES2”; “BU1”; “BU3”), hereafter non-CF #1-3, respectively (bu.edu/stemcells) (Hawkins et al., 2021; Park et al., 2017). One non-CF (“BU3,” non-CF #3) and one CF line (“C17,” Phe508del #3) had previously undergone a gene-editing strategy to insert an NKX2-1:GFP fluorescent reporter, to enable tracking and purification of lung progenitors (Crane et al., 2015; Hawkins et al., 2017). In addition, non-CF #3 also contained a TP63:TdTomato reporter used to develop the methodologies to derive airway basal cells (Hawkins et al., 2021). For each PSC line, standard quality control included G-band karyotyping and assessment of pluripotency markers (Supplementary Fig. 1); all clones had a normal karyotype.

**Figure 1.**
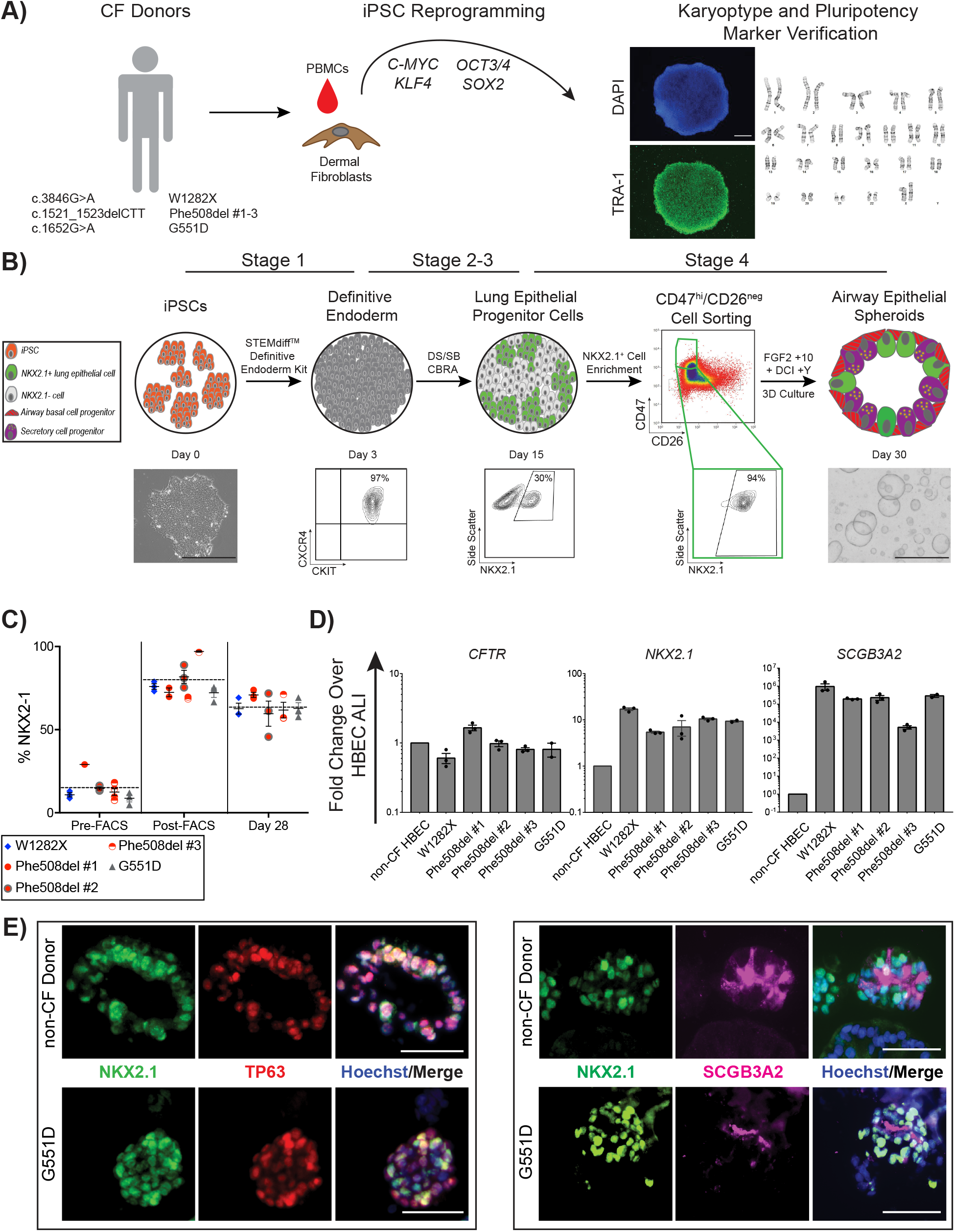
iPSCs derived from multiple CF donors are differentiated into CFTR-expressing airway epithelial cells. **A)** Schematic describing approach by which selected individuals with CF caused by different classes of mutations had somatic cells reprogrammed into iPSCs, which were subsequently tested for markers of pluripotency and G-band karyotype. Mutation details and their abbreviations are provided. An example of TRA-1 staining and DAPI nuclear labelling (scale bar represents 500µm) and normal 46XX karyotype are shown. **B)** Schematic of directed differentiation protocol to generate airway epithelial cell spheroids. Flow cytometry checkpoints shown were utilized to ensure adequate differentiation efficiency at the stages indicated. Representative images of a single iPSC colony and several airway epithelial spheroids shown (scale bars represent 500µm). **C)** Frequency of NKX2-1+ cells as a percentage of all cells at day 14-16, before (Pre-FACS), immediately after sorting CD47^hi^/CD26^neg^ cells (Post-FACS), and 14 days later (Day 28). Each point represents an individual experiment for the iPSC line indicated in the key. Horizontal dashed lines represent the average NKX2-1% across all cell lines at the designated time point. **D)** mRNA expression of canonical airway epithelial cell markers on day 28-30 of differentiation in iPSC-derived airway epithelial spheroids from the iPSC lines indicated, calculated relative to levels within non-CF HBEC derived ALI. Each point represents an individual experiment (n=3). **E)** Examples of immunolabeling of day 28-30 non-CF and CF iPSC-derived airway epithelial spheroids with antibodies against NKX2-1 and TP63 (left panels) and SCGB3A2 (right panels); scale bars represent 50µm. Lines and error bars represent mean ± standard error.

CF and non-CF PSCs were differentiated into 3-D airway epithelial spheroids precisely as previously described (Fig. 1b) (Hawkins et al., 2017; McCauley et al., 2017). The airway epithelial cell directed differentiation protocol consists of four stages that recapitulate major developmental milestones; (stage 1, day 0 to 3) definitive endoderm induction, (stage 2, day 3 to 6) anterior foregut patterning, (stage 3, day 6 to 15) lung specification, and (stage 4, day 15-30) airway patterning (McCauley et al., 2017). In an effort to achieve consistent differentiations in all 8 PSC lines a number of quality control metrics were used. For all lines we tested the efficiency of definitive endoderm induction in stage 1 by measuring the percentage of cells co-expressing the surface markers C-KIT and CXCR4 by flow cytometry (Hawkins et al., 2017; Kubo et al., 2004). All lines generated definitive endoderm with high efficiency (90.8 to 98.1% C-KIT+/CXCR4+) (Supplementary Fig. 2). To assess the efficiency of lung specification after stage 3 of the directed differentiation, we quantified the percentage of cells expressing NKX2-1 on day 14-16 by intracellular flow cytometry and similar to previous reports, observed variable NKX2-1 expression (15.1% ± 3.6%, mean ± SEM; range 4.7-29%) (Fig. 1c; Supplementary Fig. 2). To overcome this variable efficiency, we used a cell surface marker sort strategy (CD47^hi^/CD26^neg^) to enrich for NKX2-1+ cells (Fig. 1b,c) (Hawkins et al., 2017). Sorting CD47^hi^/CD26^neg^ cells significantly enriched the differentiation for NKX2-1+ cells in all lines (79.9 ± 4.6%, mean ± SEM) (Fig. 1c). In the case of the NKX2-1:GFP reporter lines (non-CF #3, Phe508del #3), NKX2-1^GFP+^ sorting was employed in place of the CD47^hi^/CD26^neg^ strategy (Supplementary Fig. 2). In stage 4 of the differentiation protocol, NKX2-1+ lung progenitors were plated in 3-D culture in a media composed of FGF2, FGF10, dexamethasone, cyclic AMP, 3-isobutyl-1-methylxanthine, and Y-27632, hereafter “FGF2+10+DCI+Y” (Fig. 1b). We previously reported that in the absence of exogenous activators of WNT signaling, NKX2-1+ lung progenitors formed epithelial spheroids composed of cells which express proximal airway markers including *SOX2, TP63* and *SCGB3A2* (McCauley et al., 2017; McCauley et al., 2018a). Expression of canonical airway epithelial cell markers in CF iPSC-derived spheroids in 3-D culture was compared to that of non-CF primary airway ALI cultures (Fig. 1d). Compared to primary cells, iPSC-derived spheroids expressed similar levels of *CFTR*. As expected based on our prior report, *NKX2-1* and *SCGB3A2* were expressed at higher levels in iPSC-derived samples compared to HBEC ALI cultures (Fig. 1d) (McCauley et al., 2017). Consistent with earlier studies we did not identify mature ciliated (*FOXJ1*) or secretory (*SCGB1A1*) cells in iPSC-derived airway spheroids at this day 28-30 time point. 63.5 ± 1.9% (mean ± SEM) of cells expressed NKX2-1 at day 28-30 (Fig. 1c). By immunolabeling, CF and non-CF spheroids were composed of NKX2-1+ cells, a subset of which co-expressed TP63 and SCGB3A2 (Fig. 1e). In summary, we generated a bank of CF iPSCs from different classes of CFTR mutations and demonstrated the differentiation of CF iPSCs into airway spheroids that express *CFTR* at similar levels to primary airway epithelial cells.

### Characterization of FIS assay in non-CF iPSC-derived airway spheroids

Through its action on adenylyl cyclase, forskolin activates cAMP-dependent channels, including CFTR (McCray et al., 1992). We first sought to determine the CFTR-dependent swelling kinetics in response to forskolin in iPSC-derived cells with normal CFTR (Fig. 2a). Using the same culture conditions as above, non-CF airway spheroids from one iPSC line (non-CF #2; n=3 experiments) were stimulated with 5µM forskolin or vehicle control (DMSO) and imaged hourly for 24 hours (Fig. 2b). Individual sphere cross-sectional surface area (CSA) was measured at each time point and calculated as a percentage of initial CSA (Fig. 2b, top panel). Within an experiment, individual spheroids variably swelled (range 112-250%), though average swelling within a well was similar between independent experiments (range 159-181%). Swelling began within 2 hours and continued until hour 20, followed by a plateau phase (20-24 hours) (Fig. 2b, middle panel). By 24 hours, forskolin-stimulated spheroid CSA increased by 173 ± 7% compared to 133 ± 4% (mean ± SEM) for vehicle treated controls (p=0.008) (Fig. 2b, bottom panel). Further FIS experiments were analyzed after 20-24 hours of forskolin exposure.

**Figure 2.**
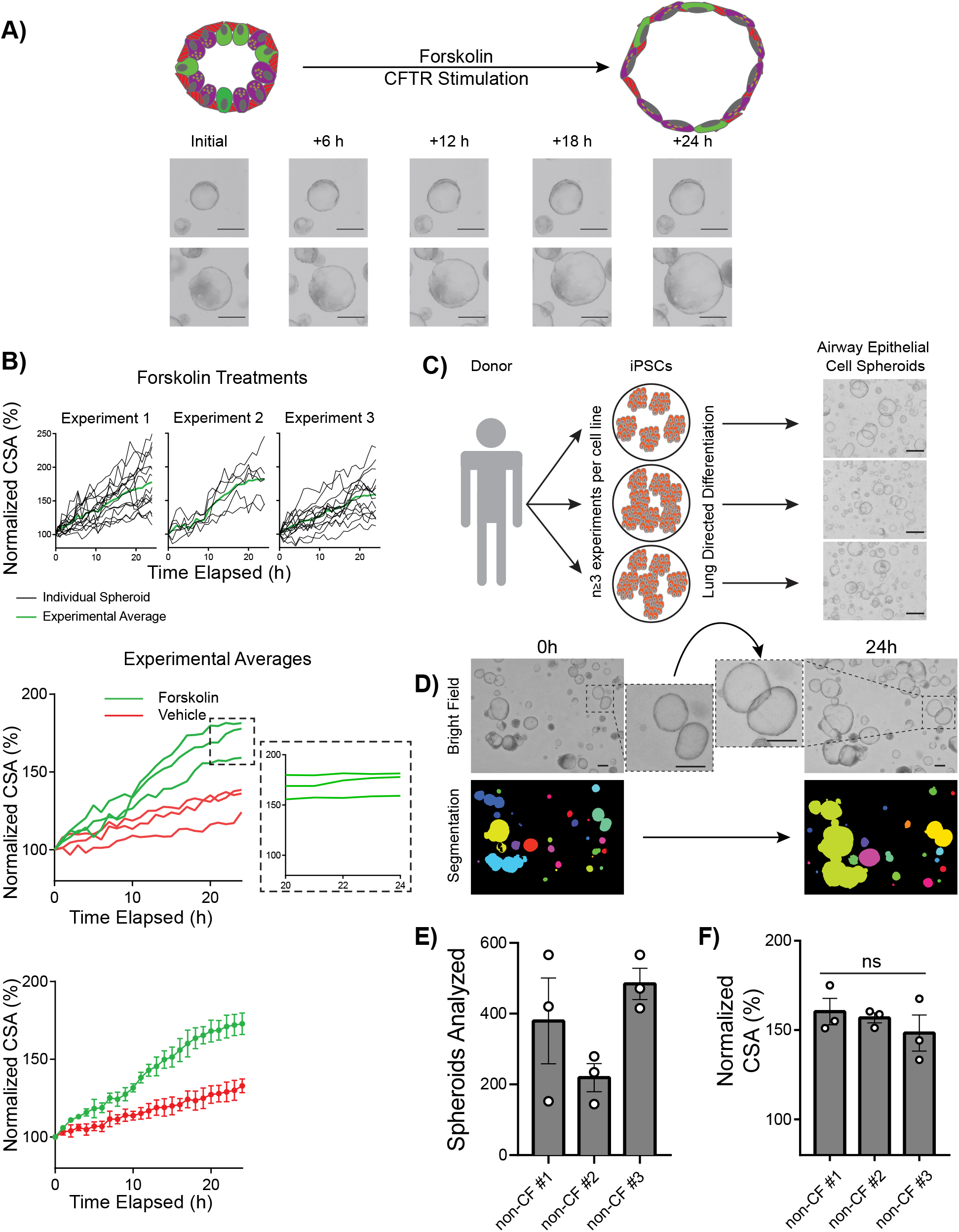
Quantification of FIS in iPSC-derived airway epithelial spheroids from non-CF donors. **A)** Schematic of FIS assay with representative images of two spheres immediately prior to as well as 6, 12, 18, and 24 hours after forskolin addition. **B)** Kinetics of non-CF airway spheroid FIS measured by change in cross-sectional area (CSA). Top panel shows individual spheroid FIS (black lines) and average FIS (green lines) (n=3 experiments). Middle panel shows compiled FIS (and vehicle control) experimental averages with magnified inset indicating 20-24h. Bottom panel shows mean and standard error for each timepoint depicted in middle panel (n=3 experiments). **C)** Experimental approach for the FIS assay. Each iPSC line was differentiated in at least three separate experiments and FIS performed on day 28-32. Examples of non-CF airway epithelial spheroids (day 28) shown on the right. **D)** Automated imaging analysis of non-CF spheroids before (left) and after (right) forskolin stimulation using OrganoSeg™ to quantify change in CSA. **E)** Number of airway spheroids analyzed per experiment for the non-CF cell line indicated. Each point represents an independent experiment. **F)** Three non-CF donor FIS responses shown. Each point represents an independent experiment Scale bars represent 250µm. Lines and error bars represent mean ± standard error.

To more broadly assess the forskolin effect between multiple non-CF genetic backgrounds (n=3 cell lines) we generated airway epithelial cell spheroids from the remaining non-CF iPSC lines (non-CF #1,3) (n=3 experiments per line) (Fig. 2c). Morphologically, non-CF spheroids were heterogeneous in size, thin-walled, and had a hollow central lumen which increased in size after forskolin treatment (Fig. 2d, insets); average cell numbers per well were similar amongst all samples (Supplementary Fig. 4). We imaged live airway epithelial cell spheroids immediately prior to, and 20-24 hours after the addition of forskolin or vehicle control (Fig. 2d). Spheroid number and CSA were calculated using automated analytical software (see Methods) (Borten et al., 2018) (Fig. 2d). For each well, we calculated the “Normalized CSA” using the equation: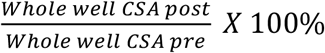 to assess the mean change in spheroid size after forskolin stimulation. We analyzed 135 ± 26 spheroids per well, 360 ± 77 spheroids for an individual experiment (mean ± SEM) (Fig. 2e), thus on average more than 1000 spheroids for each individual iPSC line. After forskolin stimulation, non-CF spheroid CSA (n=3 iPSC lines; n=3 independent differentiations) increased to 155 ± 4%, with no significant difference between genetic backgrounds, compared to 128 ± 4% for vehicle treated spheroids (p<0.0001) (Fig. 2f).

Consistent with our prior work identifying some non-lung endoderm emerging at this time-point, we observed that some cells had lost NKX2-1 expression by day 28-32 (Fig. 1c) (McCauley et al., 2018a). We assessed the contribution of non-lung (i.e. NKX2-1-) cells to the FIS response. By utilizing the NKX2-1:GFP fluorescent reporter line (non-CF #3), we identified NKX2-1-GFP+ and NKX2-1-GFP-spheroids with live cell fluorescent microscopy before and after forskolin stimulation and found no significant difference between NKX2-1+ and NKX2-1-spheroids (p=0.60) (Supplementary Fig. 3c). In summary, we developed a platform to consistently measure FIS using non-CF iPSC-derived airway epithelial spheroids from multiple genetic backgrounds.

### Genotype-dependent modulator rescue of CFTR function in a FIS assay

Having established the baseline FIS response of iPSC-derived airway spheroids from patients with a normal *CFTR* sequence, we next asked if airway spheroids derived from CF donors differed. Phe508del and G551D iPSCs were differentiated into airway spheroids and characterized as described above. Prior to forskolin stimulation, CF spheroids were smaller and appeared more dense than non-CF spheroids (Fig. 3a). Baseline spheroid size varied with the underlying *CFTR* mutation such that non-CF spheroids (0.124 ± 0.012 mm^2^) were larger than G551D (0.090 ± 0.012 mm^2^) and Phe508del spheroids (0.045 ± 0.003 mm^2^) (mean ± SEM) (Fig. 3b).

**Figure 3.**
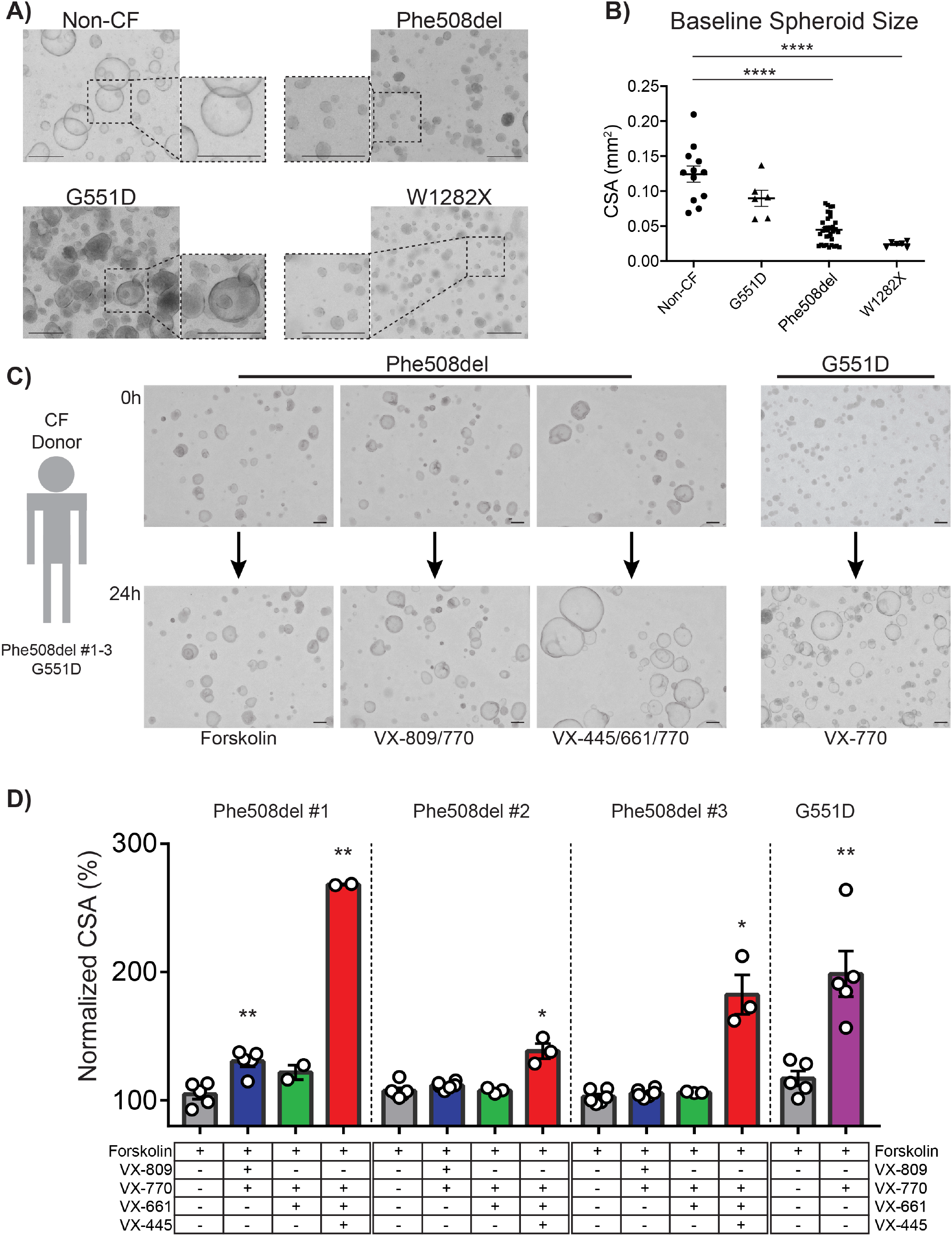
Characterization of CF iPSC-derived airway epithelial spheroids in terms of morphology, size, and FIS with and without CFTR modulator treatment. **A)** Representative microscopy images of airway spheroids from CF and non-CF donors demonstrate differences in morphologic appearance; dashed boxes depict magnified views. **B)** Baseline spheroid size differs with *CFTR* genotype. Each point represents average spheroid size before forskolin stimulation within an individual experiment (n ≥ 6 experiments per *CFTR* genotype). **C)** Representative spheroid images for the *CFTR* mutations indicated before (0h) and after (24h) treatment with compounds shown at bottom of panel. **D)** Change in CSA after treatment of CF airway epithelial cell spheroids with CFTR modulators. P-values were calculated using paired Student’s t-test comparing treatment sample to forskolin only control; * p<0.05; ** p <0.01; *** p<0.001; **** p<0.0001. Scale bars represent 250µm. Lines and error bars represents mean ± standard error.

We next tested baseline FIS of CF spheroids. We analyzed 582 ± 189 spheroids for an individual experiment (mean ± SEM) (Supplementary Fig. 4). There was small but statistically significant FIS in G551D spheroids (p=0.04) but no detectable FIS in Phe508del spheroids (Fig. 3d), consistent with their expected baseline levels of CFTR function. We next tested the effect of CFTR modulator treatment on CF spheroids (Fig. 3c). FIS after treatment of G551D spheroids with the CFTR channel potentiator VX-770 increased the CSA of spheroids by 198 ± 18% (p=0.005) (Fig. 3d). Phe508del #1-3 spheroids were treated with currently approved CFTR modulator regimens (VX-809, VX-661, VX-445/VX-661, all in combination with VX-770) (Fig. 3c,d). Treatment with first-generation correctors (i.e. VX-809, VX-661) had a small effect, significant in only 1 out of 3 patient lines (Phe508del #1, VX-809 treatment, p=0.0002). However, treatment with the recently approved VX-445 in combination with VX-661/VX-770 led to a robust increase in size of all three Phe508del patient cell lines, consistent with known *in vitro* and clinical effects of VX-445/661/770 (Fig. 3d) (187 ± 20% across three Phe508del lines).

To test the scalability of this platform for potential drug screening, we utilized an automated liquid handler to plate and treat airway epithelial spheroids using the G551D cell line in a 384-well format. Similar to the above experiments, treatment with VX-770 showed significant CSA increase (155 ± 5.7%) compared to control (111 ± 2.3%) (mean ± SEM) (p<0.0001) (Supplementary Fig. 5). Thus, we developed a platform that enables the testing of thousands of patient-derived CFTR-expressing airway spheroids across multiple classes of *CFTR* mutation with detection of baseline as well as modulator rescue of CFTR channel function. With the relative ease in generating a large number of airway epithelial cell spheroids and the scalability potential, this platform provides a new tool for patient-specific testing of drug responsiveness and for the development of novel therapeutic strategies.

### Electrophysiologic assessment of CFTR rescue in a polarized iPSC-derived airway epithelium

The gold-standard platform for the accurate assessment of CFTR function is the measurement of ion transport across differentiated primary human airway epithelium in ALI culture (Clancy et al., 2019). In our established basal cell differentiation protocol, the aforementioned Phe508del iPSC-derived airway epithelial cells (Fig. 1b, day 30) were further specified to NGFR+ airway basal cells as previously described (Fig. 4a, Supplementary Fig. 6) (Hawkins et al., 2021). After 8-14 days of ALI culture (Pneumacult ALI medium), the epithelial layer maintained barrier function (dry apical surface) and transepithelial electrical resistance (TEER) was 887 ± 189 Ω x cm^2^ (mean ± SEM) (n=3 cell lines) (Supplementary Fig. 6). We observed the presence of motile cilia and mucus hurricanes, a phenomenon similar to primary HBEC ALI cultures. When compared to non-CF HBEC ALI cultures, iPSC-derived cells expressed similar levels of relevant airway markers including *NKX2-1, SCGB1A1, FOXJ1, MUC5AC*, and *MUC5B*; as previously described, *SCGB3A2* was expressed at higher levels (Fig. 4b) (Hawkins et al., 2021). Notably, *CFTR* was expressed at similar levels to non-CF primary controls. Immunolabeling of ALI cultures confirmed the epithelium was composed of secretory (MUC5AC+), multiciliated (acetylated-α-tubulin+) and basal cells (KRT5+) (Fig. 4c).

**Figure 4.**
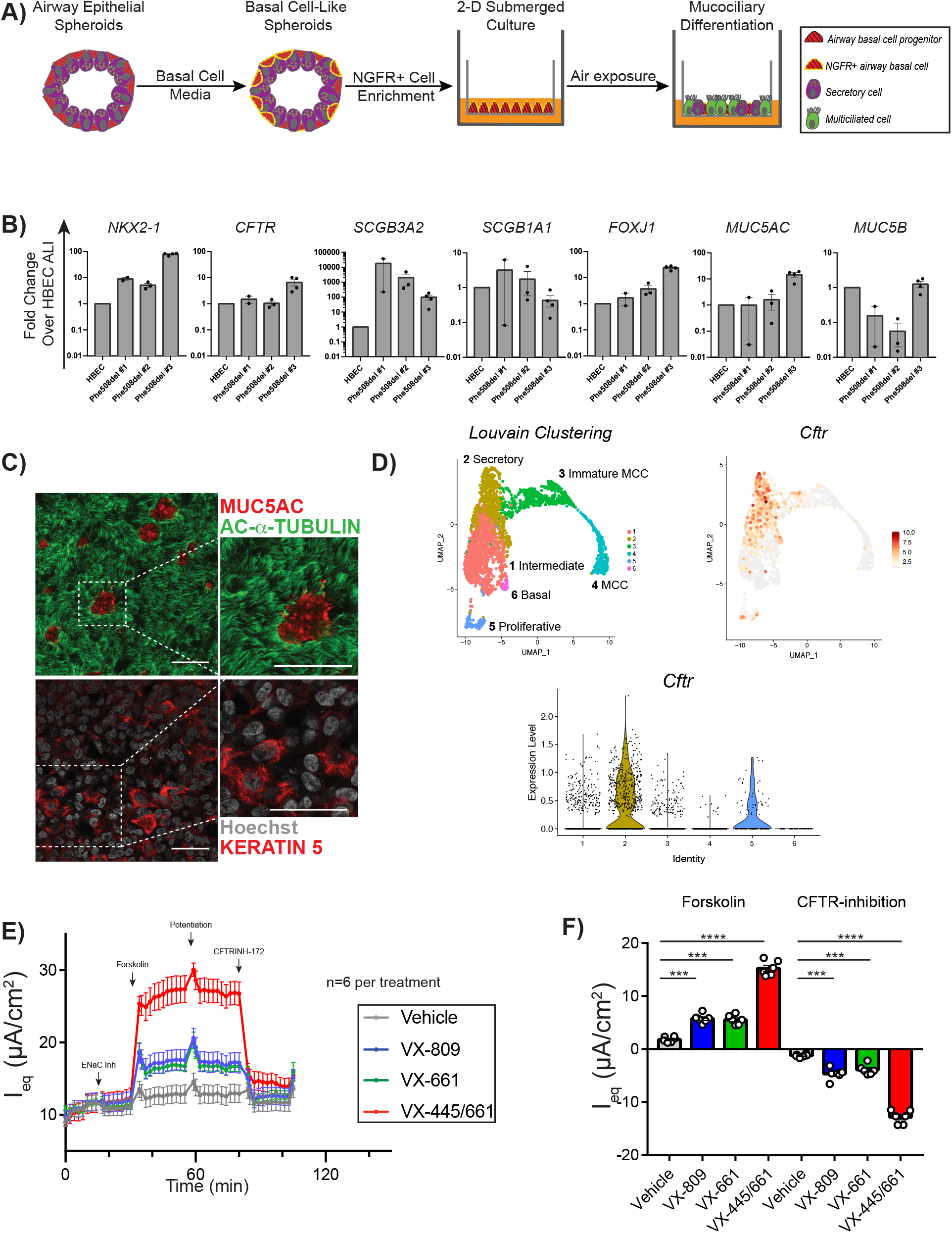
iPSC-derived airway epithelial cells grow in ALI culture and can be used to detect CFTR-dependent ion flow. **A)** Schematic of the generation of iPSC-derived ALI cultures via FACS-purified NGFR+ airway basal cells. **B)** mRNA expression of canonical airway epithelial markers within iPSC-derived ALI cultures, compared to non-CF HBEC ALI cultures. Each point represents an individual experiment. **C)** Example of immunolabeling of iPSC-derived Phe508del ALI cultures with antibodies against markers of multiciliated (acetylated-α-Tubulin), mucus secreting (MUC5AC), and airway basal cells (KRT5) with nuclei counterstained with Hoechst. **D)** Uniform Manifold Approximation and Projections (UMAPs) of scRNA-seq of non-CF iPSC-derived airway ALI cultures depicting Louvain clustering and violin plots demonstrating the expression of CFTR in the annotated clusters. **E)** Equivalent current measurements of Phe508del ALI cultures (n=6 per treatment). For both D) and E), measurements are shown at baseline and after ENaC inhibition, forskolin treatment, CFTR potentiation with Genistein, and CFTR inhibition (arrows). **F)** Quantification of peak delta forskolin and CFTR-inhibition effects of assay results shown in figure 4e. P-values were calculated using paired Student’s t-test; ***p<0.001; ****p<0.0001. Lines and error bars represent means ± standard error. Scale bars represent 50µm.

To ascertain the cell type(s) expressing *CFTR*, we re-analyzed a recent single-cell RNA sequencing (scRNA-seq) dataset from non-CF #3 iPSC-derived mucociliary cells, which were derived using a nearly identical iPSC-directed differentiation protocol (Hawkins CSC 2020). In our prior analysis, Louvain clustering identified six distinct clusters of airway epithelial cells, annotated as follows: 1) intermediate cells; 2) secretory cells; 3) immature multiciliated cells (MCC); 4) MCCs; 5) proliferative cells; 6) basal cells. We found that the majority of *CFTR* transcripts were expressed in cluster 2 (secretory), where the top DEGs included *SCGB1A1, SCGB3A2, MUC5B*, amongst others, similar to recent reports from primary airway datasets suggesting the majority of *CFTR* transcripts in human airways are in secretory cells (Carraro et al., 2020; Okuda et al., 2021).

We next tested whether Phe508del iPSC-derived ALI cultures displayed similar electrophysiologic characteristics and modulator rescue to that of published primary HBEC results. In well-differentiated iPSC-derived ALI cultures (Phe508del #1-3) with adequate TEER (see Methods), cells were pre-treated with VX-809, VX-661, VX-445/661, or vehicle control. We measured equivalent current (I_eq_) at baseline and after 1) ENaC inhibition, 2) forskolin stimulation, 3) CFTR potentiation, and 4) CFTR inhibition (Liang et al., 2017) (Fig. 4e-f). Baseline current ranged from 2.5 - 13 µA/cm^2^ without significant decrement following ENaC inhibition, similar to previous findings and suggesting low ENaC activity (Fig. 4e-f; Supplementary Fig. 7) (Hawkins et al., 2021). Treatment with CFTR modulators led to significant improvement in forskolin-stimulated current and CFTR-inhibition in all Phe508del samples, though magnitude varied between individual lines (Fig. 4f; Supplementary Fig. 6-7). Comparison between CFTR modulators showed higher CFTR-specific currents after treatment with the newest generation modulators (12.9 ± 0.5 µA/cm^2^), but again the magnitude varied between cell lines (Fig. 4f). Overall, we determined that iPSC-derived airway cells from Phe508del CF donors can generate mucociliary cultures that express *CFTR* and recapitulate the electrophysiologic baseline and pharmacologic rescue of CFTR-specific current with known modulators.

### Assessment of CFTR function in W1282X iPSC-derived airway epithelial cells

We next applied both the FIS assay of 3-D spheroids and electrophysiologic assessment of ALI cultures to measure CFTR function in W1282X iPSC-derived airway epithelia. Similar to above, we generated airway epithelial cell spheroids and iPSC-derived mucociliary cultures. In the indicated experiments, NKX2-1+ lung epithelial progenitors were purified by sorting for cells expressing the surface marker carboxypeptidase M (CPM) instead of CD47^hi^/CD26^neg^ (Gotoh et al., 2014). FIS assay of airway spheroids using approved CFTR modulators for class 2-3 mutations (VX-809, VX-661, VX-770) led to no swelling of W1282X spheroids as expected (data not shown). As no targeted treatments are currently available for individuals with class I mutations, we tested experimental compounds including a read-through agent (G418) and an inhibitor of the NMD pathway (SMG1i) in combination with CFTR modulators (Fig. 5b). Inhibition of SMG1 was recently demonstrated to increase *CFTR* mRNA and protein quantity, as well as chloride current in W1282X expressing cells (Keenan et al., 2019; Valley et al., 2019). In both iPSC-derived airway spheroids and mucociliary cultures, treatment with a SMG1i for 48 hours in the basal medium increased *CFTR* mRNA levels by 5.8 ± 0.42 and 5.5 ± 0.46 fold, respectively (mean ± SEM) (Fig. 5a). We next measured FIS of W1282X airway spheroids at baseline and in response to treatment with SMG1i, the aminoglycoside G418, and VX-445/VX-661/VX-770 to measure functional improvement in CFTR activity (Fig. 5b). Treatment with CFTR modulators alone or in combination with G418 did not lead to a significant FIS response; however, combinatorial treatment with VX-445/VX-661/VX-770, G418 and SMG1i led to a significant increase in FIS (117 ± 2% of baseline size, p = 0.03) (Fig. 5b). Finally, we generated mucociliary ALI cultures from W1282X iBCs as above. Similar to cultures generated from other healthy and CF iPSC lines, we observed an intact airway epithelium composed of basal, secretory and multicilated cells (Fig. 5c). To compare iPSC-derived cultures with primary cells, we obtained HBECs from the explanted lungs of a separate individual, also homozygous for the W1282X mutation, and subsequently generated mucociliary cultures using well-established protocols (see methods for cell line and culture details). We tested the electrophysiologic response of both iPSC- and HBEC-derived mucociliary cultures at baseline and after treatment with combinations of CFTR modulators, SMG1i and G418. Minimal ENaC-dependent current was detected in iPSC samples. In both iPSC- and HBEC-derived cultures, there were similar levels of CFTR-dependent current and patterns of responses to treatment combinations (Fig. 5d-f). In both cell types, treatment with the combination of G418, SMG1i, and modulators led to a significant improvement in peak delta forskolin current (6 ± 0.85 and 4 ± 0.2 µA/cm^2^) and CFTR-inhibited current of 4.6 ± 0.37 and 4.8 ± 0.42 µA/cm^2^ respectively, with no significant differences between iPSC- and HBEC-derived results (Fig. 5d-f).

**Figure 5.**
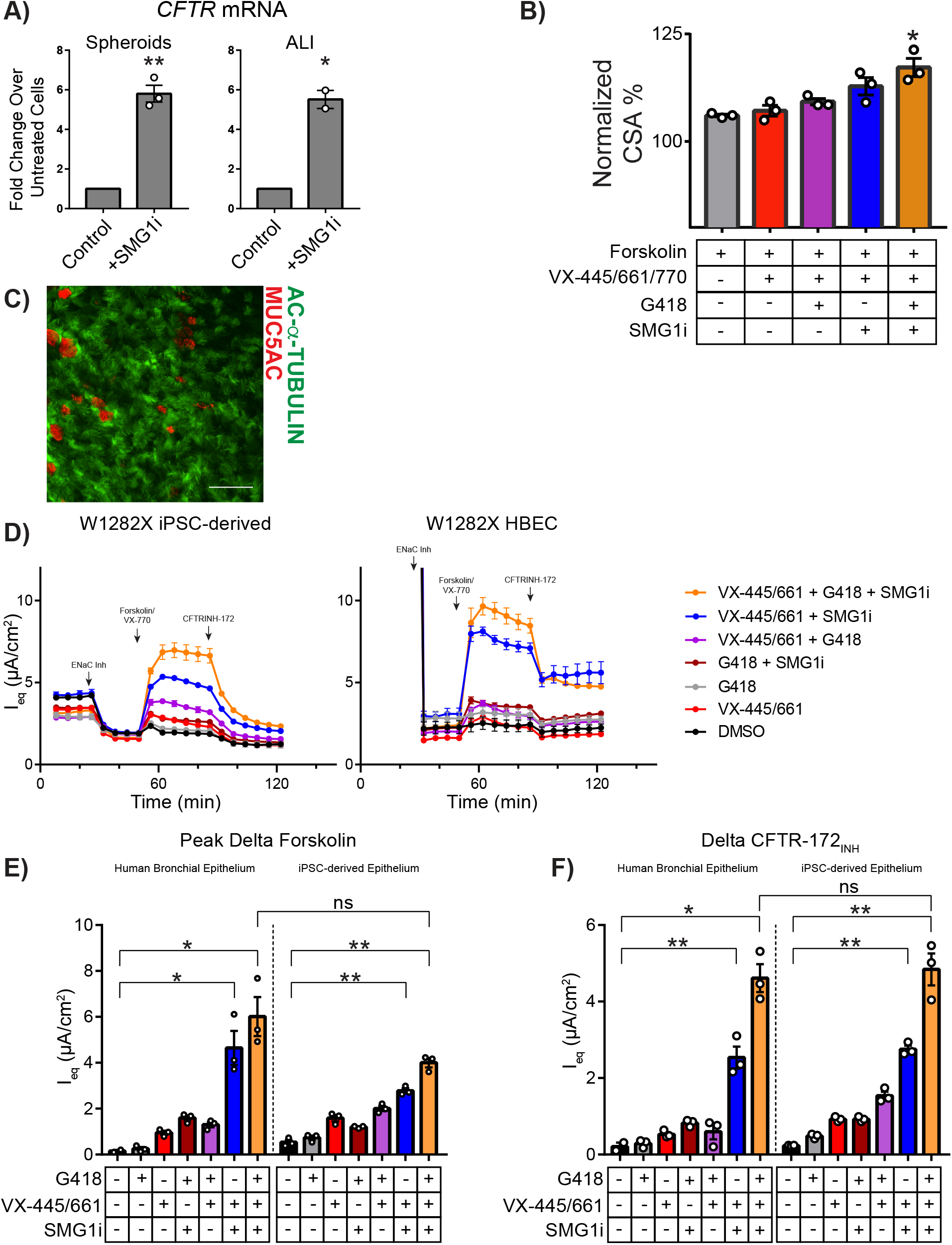
Assessment of CFTR function in W1282X iPSC-derived airway spheroids and mucociliary cultures. **A)** Treatment with SMG1i increases mRNA expression levels of *CFTR* in W1282X iPSC-derived spheroids (left) and mucociliary ALI cultures (right) compared to controls. **B)** FIS assay of W1282X iPSC-derived airway spheroids after combination treatment with VX-445/VX-661/VX-770, G418, and SMG1i (n=3). **C)** Example of immunolabeling of iPSC-derived W1282X ALI cultures with antibodies against markers of multiciliated (acetylated-α-Tubulin) and mucus secreting (MUC5AC) cells. **D)** Electrophysiological assessment of W1282X iPSC-derived (left) and primary mucociliary cells (right) using a TECC24 instrument after the indicated treatments. **E-F)** Quantification of equivalent current responses from curves in figure 5D, depicting the peak forskolin-generated (E) and CFTR-inhibited (F) currents after the treatments indicated. P-values were calculated using paired Student’s t-test; * p<0.05; ** p <0.01; *** p<0.001; **** p<0.0001. Scale bars represent 50µm. Lines and error bars represents mean ± standard error.

## Discussion

Utilizing iPSCs from individuals with class 1-3 *CFTR* mutations, we have generated *CFTR*-expressing airway epithelial cells and measured baseline as well as rescue of CFTR channel function with approved and experimental therapeutics. We adapted two established assays of CFTR function – FIS of 3-D spheroids and equivalent current measurement of polarized mucociliary epithelia. In both assays, we measured CFTR function and detected modulator rescue of CFTR function in class 2-3 mutations. Additionally, we have used these platforms to demonstrate the feasibility of identifying and developing potential novel agents for class 1 CFTR mutations such as W1282X. To our knowledge, this is the first report of an iPSC-based airway model that can detect genotype-specific CFTR modulator response. The iPSC-based system has a number of key advantages compared to current *in vitro* models including the ability to produce large numbers of disease-relevant *CFTR*-expressing airway cells from individuals with rare mutations without the need for an invasive biopsy. We anticipate that this platform will be a useful tool in the pursuit of novel therapeutic approaches for those individuals with CF who continue to struggle without targeted treatment.

The platform developed here builds on the field’s recent progress in the directed differentiation of iPSCs into functional *CFTR*-expressing airway epithelial cells (Crane et al., 2015; Firth et al., 2014; Hawkins et al., 2017; 2021; McCauley et al., 2017; Wong et al., 2012). First, we expanded on prior proof-of-concept studies using iPSC-derived airway spheroids, composed of immature basal-like and secretory-like cells, which established the CFTR-dependent swelling response to forskolin (McCauley et al., 2017). We determined the FIS kinetics in cell lines with normal CFTR and then measured the FIS in Class 1-3 CF iPSC-derived cells. This overall approach is reminiscent of the initial description of FIS in 1992, when McCray et al. first reported the CFTR-dependent swelling of fetal lung explants in response to cAMP activation using forskolin (McCray et al., 1992). More recently, this approach was comprehensively and elegantly applied to primary rectal organoids (Berkers et al., 2019; Dekkers et al., 2013; 2016). One of the major advantages of the rectal organoid system compared to other *in vitro* platforms is its potential scalability. Similar to the published rectal organoid data, iPSC-derived airway epithelial spheroids respond to forskolin in a CFTR-dependent manner and also demonstrate pharmacologic rescue of FIS in class 2 and 3 CF spheroids after treatment with CFTR modulators (Dekkers et al., 2013). While we detected significant swelling in non-CF spheroids by two hours, this is notably slower than rectal organoids which swell more rapidly and to a greater size, likely due to higher CFTR expression in rectal epithelial cells (Dekkers et al., 2013). The lower CFTR expression of airway spheroids may also explain the low FIS in response to first-generation correctors (VX-809 or VX-661) in Phe508del samples, however these molecules have relatively small clinical effects (Keating et al., 2018; Rowe et al., 2017; Wainwright et al., 2015). iPSCs may offer advantages over rectal organoids including the ability to maintain iPSC-derived *CFTR*-expressing cells in culture for extended periods. Additionally, patient derived airway epithelial cells may offer a more appropriate *in vitro* surrogate given the predominant pulmonary manifestations of CF disease. It remains unclear whether primary rectal tissue that displays no known CF pathology is suitable for *in vitro* CFTR modulator testing (Clancy et al., 2019). We also demonstrated the feasibility of scaling up to a 384-well format for drug screening experiments (Supplementary Fig. 5). Future studies will be required to determine whether the FIS of airway spheroids reflects individual disease severity or is predictive of clinical efficacy of CFTR modulators, as others have done with the rectal organoid FIS assay (Berkers et al., 2019; Dekkers et al., 2016).

Electrophysiologic measurement of CFTR-dependent ion flux in CF HBEC cultures is a fundamental tool in CF research. However, access to primary HBECs from individuals with rare mutations, who are a major research priority, is a significant bottleneck. We sought to determine the fidelity of iPSC-derived mucociliary airway epithelial cultures and found that iPSC-derived airway epithelial cultures exhibited CFTR-dependent currents and pharmacologic rescue at magnitudes comparable to primary HBEC cultures (Capurro et al., 2021; Gentzsch et al., 2017). For example, Gentzsch et al. measured CFTR-dependent currents of 3-7.5 µA/cm^2^ after treating homozygous Phe508del HBECs with VX-809 (n=3 donors) and Capurro et al. recorded 10-15 µA/cm^2^ CFTR-dependent currents in Phe508del HBECs after VX-445 treatment (n=5 donors). In iPSC-derived airway cells that generated well-differentiated mucociliary epithelia, VX-809 and VX-445/VX-661 treatment improved CFTR-specific current by 1.3-4.7 and 1.4-12.9 µA/cm^2^, respectively. We noted variability amongst the three different Phe508del iPSC lines and further work will be required to determine if there are donor-specific differences. It should be noted that several quality control steps were included in our differentiation protocol and only iPSC-derived ALI cultures with confirmed robust mucociliary differentiation and acceptable TEER measurements were included for these electrophysiological studies. Despite this and for unclear reasons, iPSC-derived cultures displayed a number of differences including minimal ENaC-dependent current and reduced durability (Supp Fig 7 for raw tracings). Since passage number and culture media can affect the cellular composition of primary airway epithelial cultures, including CFTR-expressing cell types, further work will also be focused on identifying optimal culture conditions for consistent electrophysiological assessments of CFTR function in iPSC derived cultures (Gentzsch et al., 2017). Similar to the FIS of airway organoids, determining the extent to which electrophysiologic measurements in iPSC-derived airway cultures predicts disease severity or clinical efficacy of CFTR modulators will be required. While primary HBECs remain the gold-standard assay for CF, iPSC-derived airway cells may accelerate research by overcoming the bottleneck in access to human cells carrying rare CFTR mutations with ultimate validation of candidate therapeutics in HBECs.

In a proof-of-concept study to assess the potential of iPSC-derived airway cells to detect small changes in CFTR dependent current in class 1 mutations we compared mucociliary cultures generated from iPSCs and HBECs, from separate donors homozygous for the W1282X mutation. We identified: 1) small but significant CFTR restoration with combinatorial treatment including G418, SMG1i, and CFTR modulators and 2) an overall similar magnitude of electrophysiologic response between iPSC- and HBEC-derived cultures. We acknowledge the small nature of this study with only two individuals, however, the rarity of primary W1282X HBECs is a major limiting factor in these experiments. Future efforts, heavily focusing on comparing iPSC-derived and primary airway epithelial cells from the same genotype (and donor) will be required.

This technology has several potential applications. The ability to derive an array of cell types from different organs affected by CF from iPSCs may facilitate studies focused on the multisystem nature of CF. For example, in addition to the airway cells described here, CFTR-expressing intestinal epithelial cells (Mithal et al., 2020), pancreatic exocrine cells (Simsek et al., 2016) and cholangiocytes have been described (Ogawa et al., 2015). The iPSC technology will hopefully lead to biobanks of cells from patients with CFTR mutations, particularly rare mutations, that are readily available to researchers. Since iPSC-derived airway basal cells can be efficiently cryopreserved for long-term storage while retaining their capacity to form CFTR-expressing airway epithelium in established protocols, thus the platform lends itself to creating biobanks of CF iPSC-derived airway basal cells that can be shared with the worldwide research community (Hawkins et al., 2021). The ambitious approaches aimed at treating or even curing individuals with class 1 mutations include complex pharmacotherapy, gene-editing/delivery or cell-based therapies and will require ready access to cellular platforms. Ongoing research in molecular therapeutics includes targeting NMD inhibition (as partially shown here), read-through of premature termination codons, development of novel CFTR modulators, synthetic transfer RNAs, and modified CFTR mRNAs, all of which may be tested using iPSC-derived airway cells (Pranke et al., 2019). *In vivo* gene-correction has the potential to cure CF but requires multi-disciplinary expertise to accomplish delivery of efficacious but safe gene-editing machinery to the cell-type(s) of interest in the airway epithelium. Cell-based therapy for the lungs using either autologous primary or iPSC-derived airway cells is in its infancy, but also has the theoretical potential to cure CF (Berical et al., 2019). We anticipate that the iPSC platform described here will be a useful tool in advancing these endeavors.

We note a number of limitations to the iPSC technology that will require further work. The airway differentiation protocol (like many other iPSC protocols) is lengthy and at each patterning stage we detect variability. To offset this, we have developed surface marker approaches to isolate first the lung progenitor cells of interest and later iPSC-derived airway basal cells. The reproducibility of this approach is demonstrated through the successful generation of airway epithelial cells in 8/8 cell lines analyzed in this manuscript. While non-lung endodermal cells are detected in airway spheroid cultures, we concluded that the FIS assay was not affected adversely by their presence. We anticipate that directed differentiation protocols will be further refined in time with a better understanding of the early stages of human lung development.

In conclusion, we describe here a platform using iPSC-derived airway epithelial cells that enables the detection of baseline CFTR function and can measure the degree of CFTR rescue by modulator compounds in two distinct assays. The future of CF treatment, particularly for individuals with less common *CFTR* mutations, will depend on readily available patient-derived *CFTR*-expressing cells for preclinical testing; this iPSC platform offers several potential advantages that will complement existing cell and animal models of CF.

## Acknowledgements

We thank Anne Hinds for technical support with embedding and sectioning of spheroids; Brian Tilton for expertise and technical support with cell-sorting; Sam Gallant for technical support in electrophysiology. We are indebted to Greg Miller and Marianne Janes for overall laboratory support as well as reprogramming and characterization of iPSC lines. We sincerely thank Darrell Kotton for his guidance, support, and vision of this project and research center. A.B. was supported by T32:HL7035-44, CFF BERICA2010; J.A.L. was supported by T32:HL7035-44; F.J.H. was supported by R01 HL139799-04, CFF HAWKIN20XX2 and Emily’s Entourage. S.H.R. was supported by NIH grant DK065988, CFF grants BOUCHE19RO, and RANDEL20XX2, and Emily’s Entourage. All authors critically evaluated and approved of the manuscript.

## Author Contributions

A.B. and F.J.H. conceived the work, designed the experiments, analyzed the data, and wrote the manuscript. A.B., J.A.L., M.L.B., J.L., A.M., A.S., and M.P. performed differentiation experiments. R.L., J.L., J.H., K.C., J.M., and S.R. performed the electrophysiology experiments and provided valuable discussion. N.R. and D.T. performed the differentiations and experiments utilizing the automated pipettor. G.M., K.H. and P.M. provided valuable discussion and patient samples.

## Supplementary Figure Legends

**Supplementary Figure 1. Description and characterization of pluripotent stem cell lines**. Cell lines shown with de-identified coding, as well as original cell source and *CFTR* genotype. Karyotypes were confirmed as normal for all cell lines and markers of pluripotency were assessed with immunofluorescent staining or flow cytometry as indicated. Scale bars represent 500µm.

**Supplementary Figure 2. Flow cytometry assessment of lung-directed differentiation checkpoints for CF cell lines**. Each cell line, shown in a separate row, was analyzed at the timepoints shown to ensure adequate progress of directed differentiation. Day 3 samples (column 1) were assessed by co-expression of cell surface markers CKIT and CXCR4; stained samples shown in red, isotype controls shown in blue. Samples were assessed on day 14-16 (prior to sorting), immediately post-sorting, and on day 28-30 for intracellular NKX2-1 expression as shown (columns 2, 4, 5). Samples were enriched for NKX2-1 on day 14-16 using the CD47^hi^/CD26^neg^ cell sorting strategy, shown in column 3. Flow cytometry and sorting of Phe508del #3 (row 5) utilized an NKX2-1:GFP fluorescent reporter, rather than intracellular NKX2-1 staining or CD47hi/CD26neg sorting, respectively. Numbers overlaid onto flow cytometry plots represent the percentage of cells for the gate shown. Gating was determined using known negative cells, of a similar differentiation day.

**Supplementary Figure 3. Quantification of spheroid numbers and magnitude of NKX2-1+ versus NKX2-1-spheroid FIS. A)** Spheroid numbers shown represent the number analyzed in FIS assay (note: spheroids on the edge of imaging or under the size threshold were not counted). **B)** CF and non-CF spheroids contain similar number of cells. Each point represents an individual experiment. **C)** Spheroids from non-CF #3 were generated and imaged using live cell fluorescence microscopy. Utilizing the NKX2-1:GFP fluorescent reporter, we identifed NKX2-1^GFP+^ and NKX2-1^GFP-^ spheroids prior to FIS (left panel). After 24 hours of forskolin stimulation, spheroid change in CSA is shown; each point represents an individual spheroid. Lines and error bars represent mean and standard error; scale bar represents 100µm.

**Supplementary Figure 4. Spheroid size and morphology before and after FIS**. Representative images of spheroids from indicated *CFTR* genotypes before and after forskolin stimulation. Spheroids were treated control DMSO, forskolin alone, or forskolin in combination with the treatments indicated. Scale bars represent 500µm.

**Supplementary Figure 5. iPSC-derived airway epithelial spheroids can be plated in 384-well format with demonstration of drug response**. Day 30 iPSC-derived airway epithelial spheroids were generated from G551D and plated into 384-well tissue culture plates using an automated liquid handler. CSA was compared pre- and post-treatment with quantification shown at the right. Each point represents an individual sphere. P-values shown were calculated using unpaired Student’s t-test; **** p<0.0001.

**Supplementary Figure 6. Generation and equivalent current assay of Phe508del iPSC-derived mucociliary cultures. A)** NGFR+ airway basal cells were identified using flow cytometry and sorted using the gating strategy shown. Stained samples are shown in red, isotype controls shown in blue. **B)** TEER of mucociliary cultures of Phe508del #1-3. Each point indicates an individual transwell insert. **C-D)** Compiled plots from equivalent current assays of Phe508del #1 and #2. Cultures were pre-treated with either DMSO (grey), VX-809 (blue), or VX-445/661 (red) prior to electrophysiologic assessment. Points and error bars represent mean and standard error. Shown at bottom panel is the quantification of peak forskolin and CFTR-inhibitor effects. P-values shown were calculated using unpaired Student’s t-test; * p<0.05; ** p <0.01; *** p<0.001; **** p<0.0001.

**Supplementary Figure 7. Representative raw equivalent current plots for Phe508del cell lines**. With the pre-treatments indicated, mucociliary cultures underwent electrophysiologic assessment. Four arrowheads shown indicate treatment with 1) Benzamil, 2) Forskolin, 3) Genistein, and 4) CFTR_INH_-172.

