## Supplementary Figures for "A multimodal iPSC platform for cystic fibrosis drug testing"

**Supplementary Figure 1**

| Patient Code Cell Line | CFTR Genotype | Cell Source | Karyotype | Previous References (if applicable) |
| --- | --- | --- | --- | --- |
| non-CF #1 RUES2 | Normal | Embryonic | 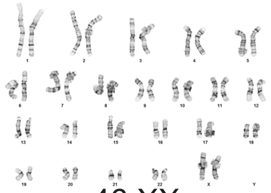 46 XX | |
| non-CF #2 BU1 | Normal | Dermal Fibroblasts | 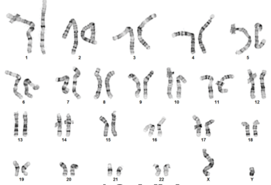 46 XY | Park et al, 2016 Mithal et al, 2020 |
| non-CF #3 BU3 | Normal | PBMCs | 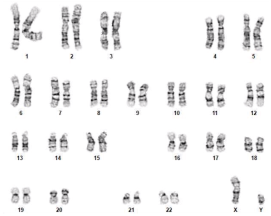 46 XY | Hawkins et al, 2021 |
| W1282X 20801 | W1282X/ W1282X | PBMCs | 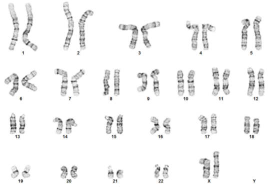 46 XX | |
| Phe508del #1 CFTR4-2 | Phe508del/ Phe508del | PBMCs | 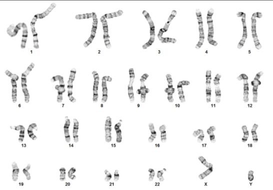 46 XY | |
| Phe508del #2 RC2 204 | Phe508del/ Phe508del | Dermal Fibroblasts | 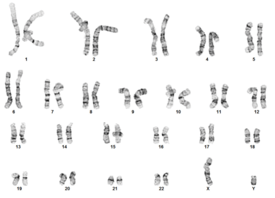 46 XY | Somers et al, 2010 |
| Phe508del #3 C17 | Phe508del/ Ile507del | Dermal Fibroblasts | 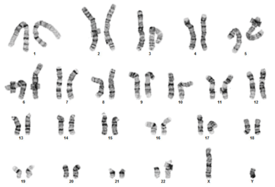 46 XY | Crane et al, 2015 |
| G551D CFTR5-5 | G551D/ G551D | PBMCs | 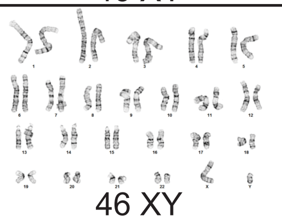 46 XY | |

### Supplementary Figure 2

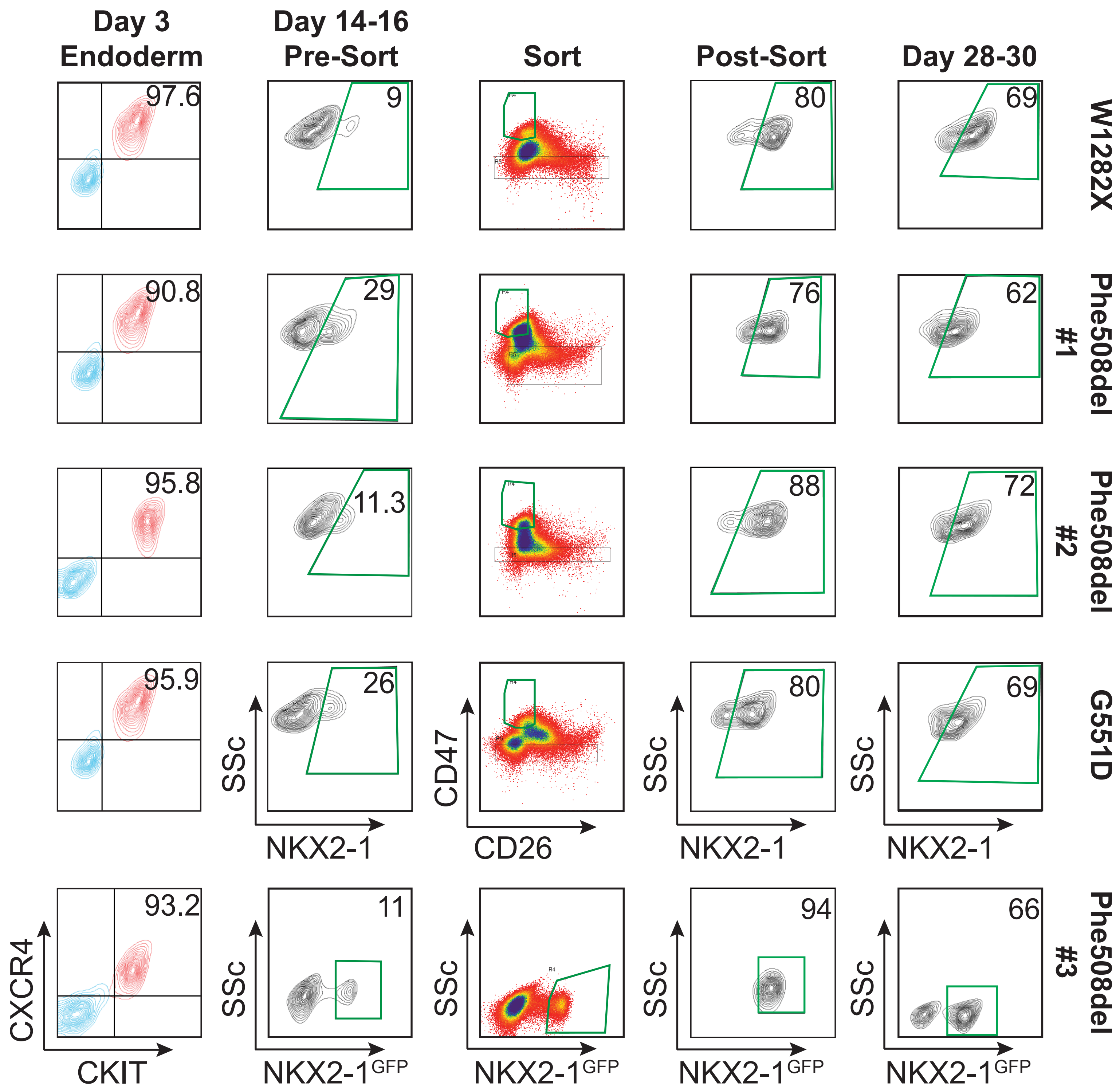

### Supplementary Figure 3

A)

| | Cell Line | Total/Experiment | Total/Condition | Experiments |
|---|---|---|---|---|
| Non-CF | non-CF #1 | 379 | 95 | 3 |
| | non-CF #2 | 219 | 110 | 3 |
| | non-CF #3 | 484 | 242 | 3 |
| CF | W1282X | 427 | 107 | 3 |
| | Phe508del #1 | 320 | 80 | 5 |
| | Phe508del #2 | 489 | 98 | 6 |
| | Phe508del #3 | 1329 | 266 | 6 |
| | G551D | 344 | 86 | 6 |
| Means | Overall | 377 | 138 | |
| | Non-CF | 361 | 149 | |
| | CF | 582 | 127 | |

B)

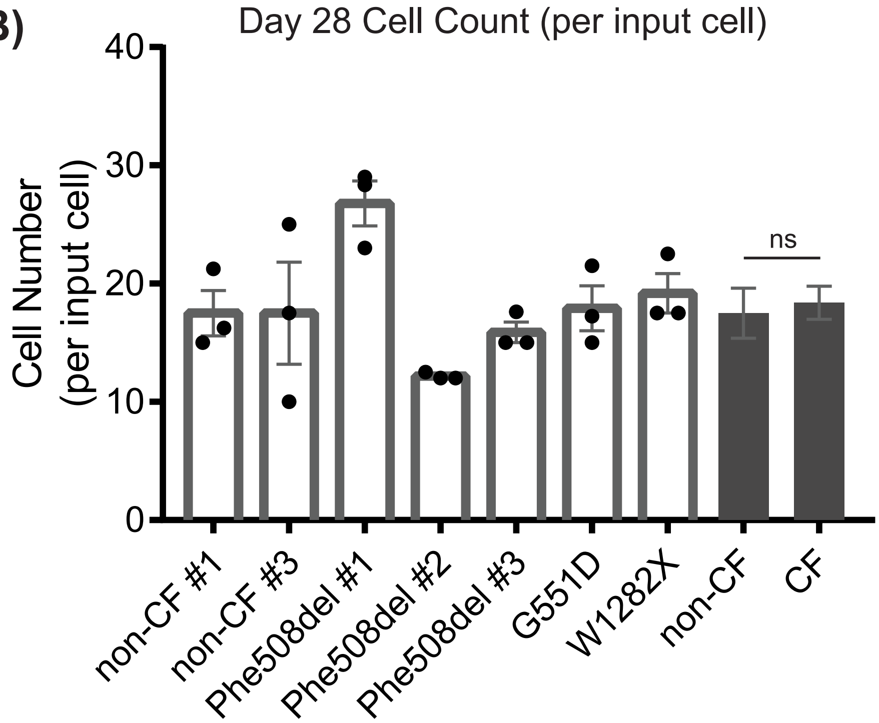


C)

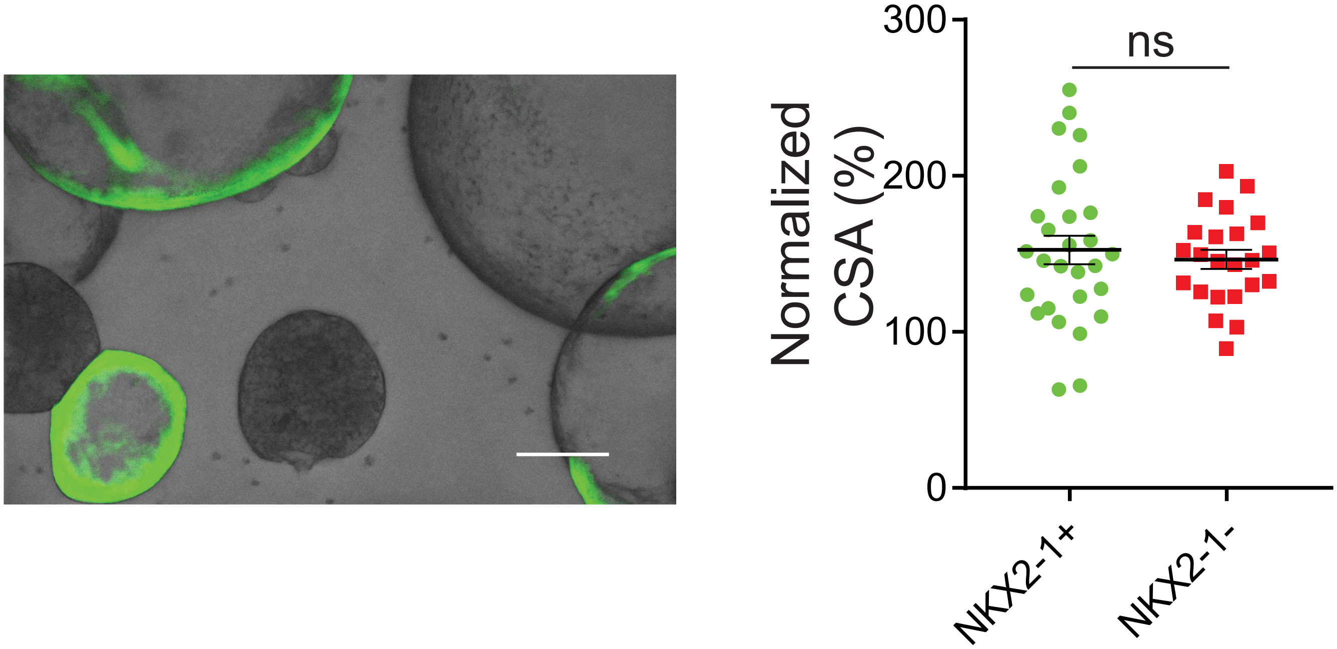

Supplementary Figure 4

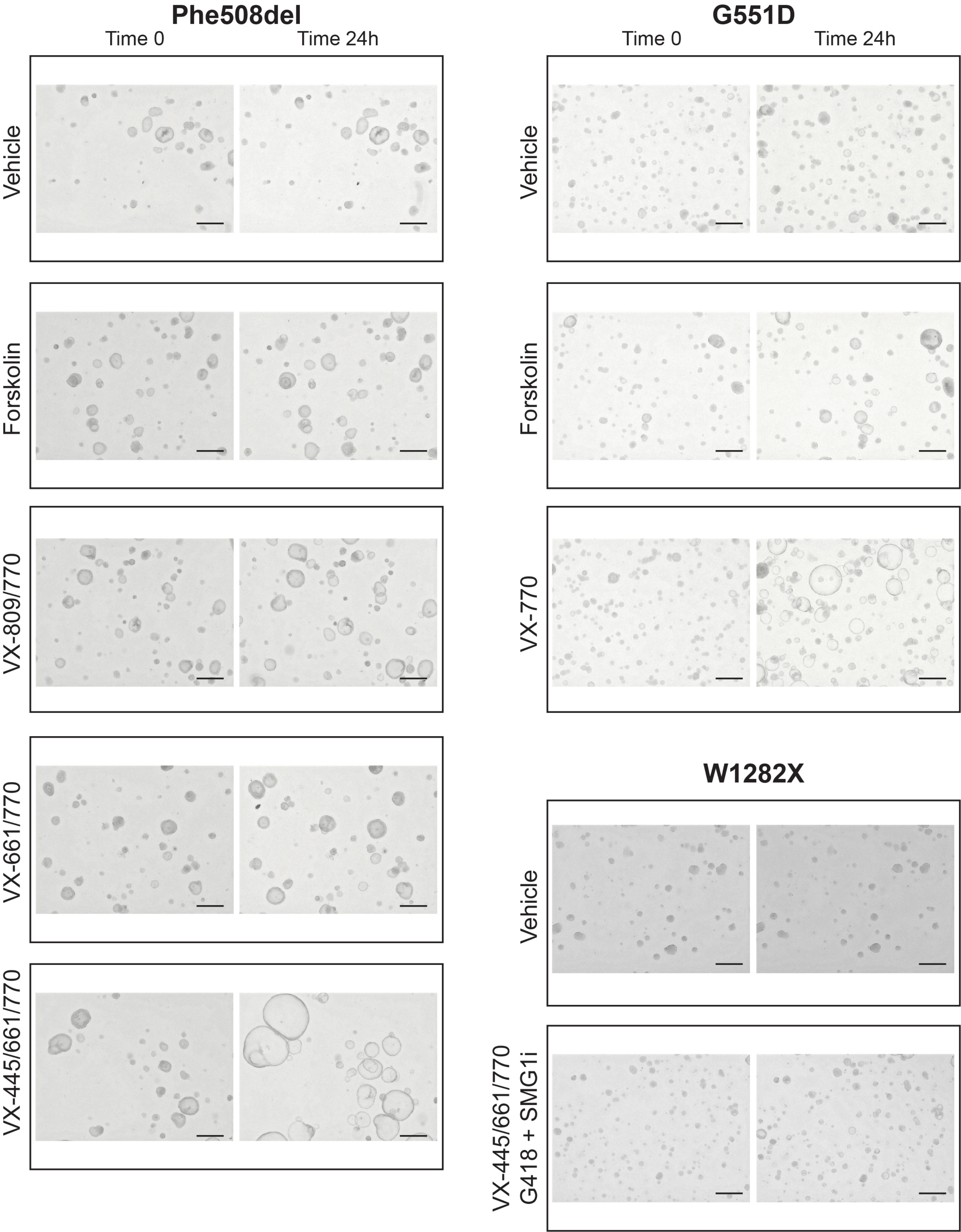

**Supplementary Figure 5**

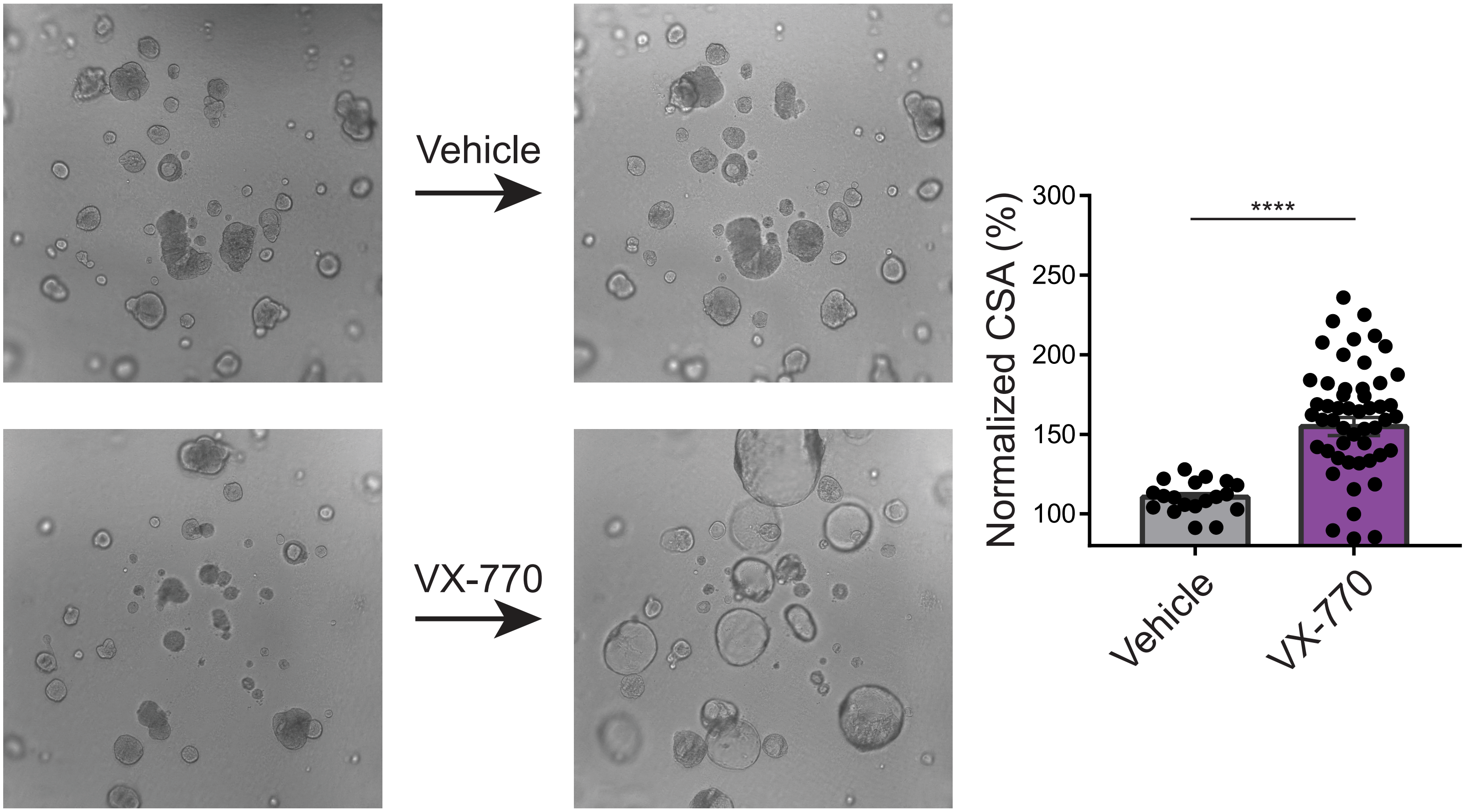

### Supplementary Figure 6

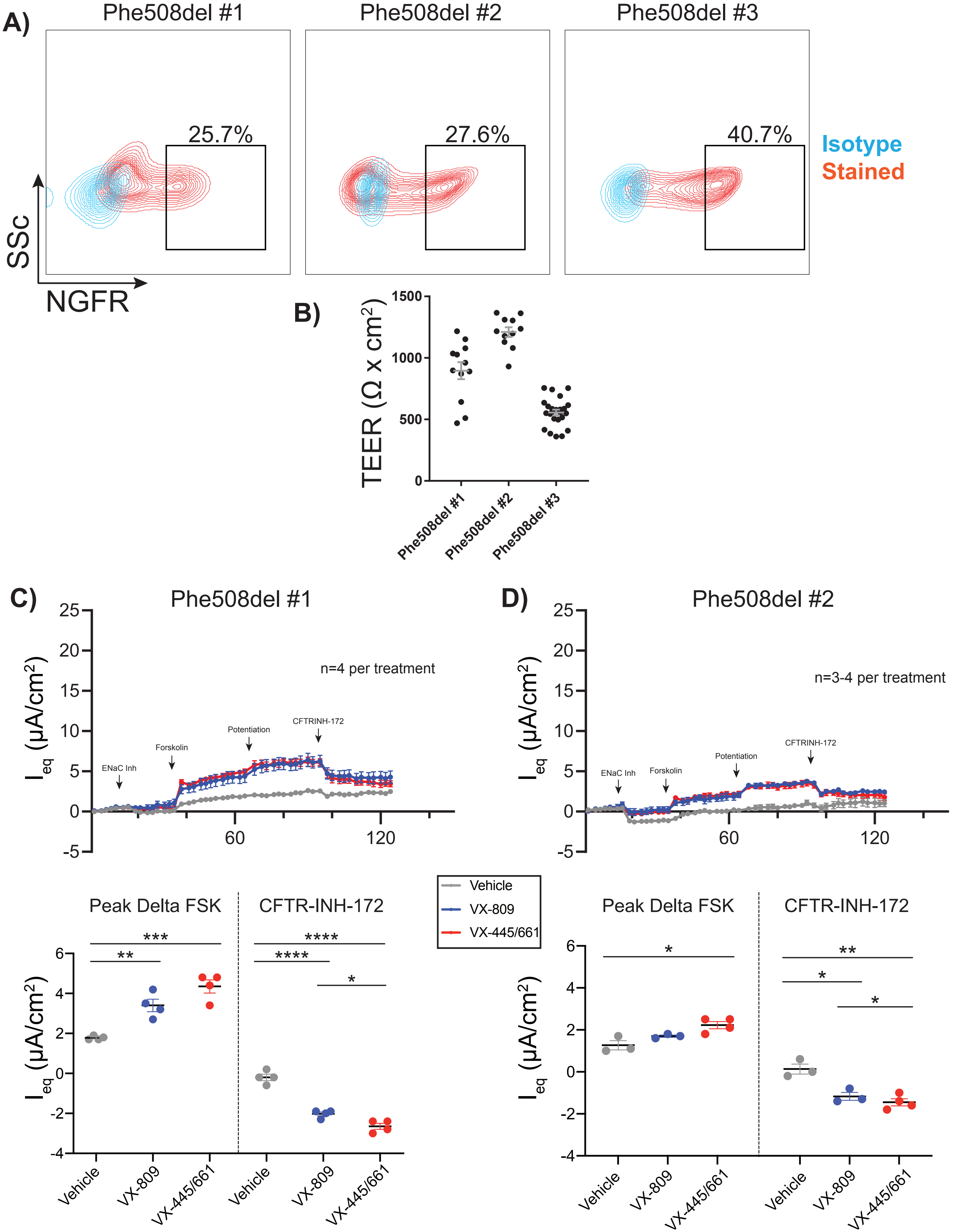

**Supplementary Figure 7**

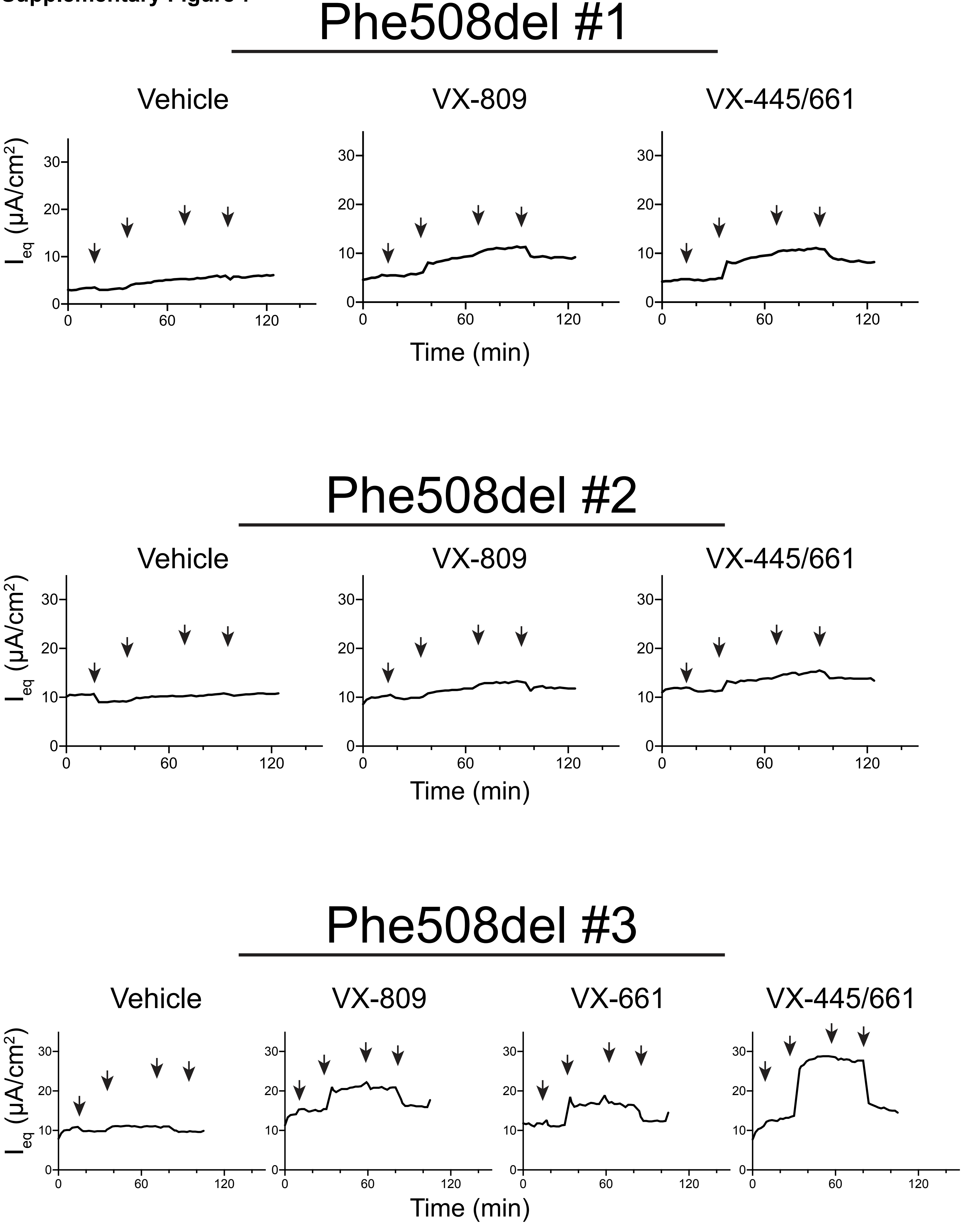
